# Anti-SARS-CoV-2 IgG from severely ill COVID-19 patients promotes macrophage hyper-inflammatory responses

**DOI:** 10.1101/2020.07.13.190140

**Authors:** Willianne Hoepel, Hung-Jen Chen, Sona Allahverdiyeva, Xue Manz, Jurjan Aman, Amsterdam UMC COVID-19 Biobank, Peter Bonta, Philip Brouwer, Steven de Taeye, Tom Caniels, Karlijn van der Straten, Korneliusz Golebski, Guillermo Griffith, René Jonkers, Mads Larsen, Federica Linty, Annette Neele, Jan Nouta, Frank van Baarle, Cornelis van Drunen, Alexander Vlaar, Godelieve de Bree, Rogier Sanders, Lisa Willemsen, Manfred Wuhrer, Harm Jan Bogaard, Marit van Gils, Gestur Vidarsson, Menno de Winther, Jeroen den Dunnen

**Affiliations:** Department of Rheumatology and Clinical Immunology, Amsterdam UMC, Amsterdam Rheumatology and Immunology Center, Amsterdam, The Netherlands; Department of Experimental Immunology, Amsterdam UMC, University of Amsterdam, Amsterdam Infection and Immunity Institute, Amsterdam, The Netherlands; Department of Medical Biochemistry, Experimental Vascular Biology, Amsterdam Cardiovascular Sciences, Amsterdam Infection and Immunity, Amsterdam UMC, University of Amsterdam, Amsterdam, The Netherlands; Department of Medical Microbiology, Amsterdam UMC, University of Amsterdam, Amsterdam Infection and Immunity Institute, Amsterdam, The Netherlands; Department of Pulmonary Medicine, Amsterdam UMC, location VUMC, Amsterdam, The Netherlands; Department of Pulmonology, Amsterdam UMC, University of Amsterdam, Amsterdam, The Netherlands; Department of Experimental Immunohematology, Sanquin Research, Amsterdam, The Netherlands, and Landsteiner Laboratory, Amsterdam UMC, University of Amsterdam, Amsterdam, The Netherlands; Department of Internal Medicine, Amsterdam UMC, University of Amsterdam, Amsterdam Infection and Immunity Institute, Amsterdam, The Netherlands; Department of Respiratory Medicine, Amsterdam UMC, University of Amsterdam, Amsterdam, The Netherlands; Center for Proteomics and Metabolomics, Leiden University Medical Center, Leiden, The Netherlands; Department of Intensive Care Medicine, Amsterdam UMC, University of Amsterdam, Amsterdam, The Netherlands; Department of Otorhinolaryngology, Amsterdam UMC, University of Amsterdam, Amsterdam

## Abstract

For yet unknown reasons, severely ill COVID-19 patients often become critically ill around the time of activation of adaptive immunity. Here, we show that anti-Spike IgG from serum of severely ill COVID-19 patients induces a hyper-inflammatory response by human macrophages, which subsequently breaks pulmonary endothelial barrier integrity and induces microvascular thrombosis. The excessive inflammatory capacity of this anti-Spike IgG is related to glycosylation changes in the IgG Fc tail. Moreover, the hyper-inflammatory response induced by anti-Spike IgG can be specifically counteracted in vitro by use of the active component of fostamatinib, an FDA- and EMA-approved therapeutic small molecule inhibitor of Syk.

**One sentence summary:** Anti-Spike IgG promotes hyper-inflammation.

## Main text

Coronavirus disease 2019 (COVID-19), which is caused by severe acute respiratory syndrome coronavirus 2 (SARS-CoV-2), is characterized by mild flu-like symptoms in the majority of patients(*1, 2*). However, approximately 20% of the cases have more severe disease outcomes, with bilateral pneumonia that may rapidly deteriorate into acute respiratory distress syndrome (ARDS) and even death by respiratory failure. With high numbers of people being infected worldwide and very limited treatments available, safe and effective therapies for the most severe cases of COVID-19 are urgently needed.

Remarkably, many of the COVID-19 patients with severe disease show a dramatic worsening of the disease around 1-2 weeks after onset of symptoms(*2, 3*). This is suggested not to be a direct effect of viral infection, but instead to be caused by over-activation of the immune system, particularly because it coincides with the activation of adaptive immunity(*2*). This excessive immune response is frequently described as a ‘cytokine storm’, characterized by high levels of pro-inflammatory cytokines(*3, 4*). Detailed assessment of the cytokine profile in severe cases of COVID-19 indicates that some cytokines and chemokines are particularly elevated, such as IL-6, IL-8, and TNF(*5–7*). In contrast, type I and III interferon (IFN) responses, which are critical for (early phase) anti-viral immunity, appear to be suppressed(*8*). Combined, the high pro-inflammatory cytokines, known to induce collateral damage to tissues, together with muted anti-viral responses, indicate a highly unfavorable immune response in severe cases of COVID-19.

Previously, it has been shown that the virus leading to SARS, SARS-CoV, causes severe inflammation and lung injury through IgG antibodies(*9*). Using a SARS-CoV macaque model, Liu et al. demonstrated that the early emergence of anti-CoV Spike-protein IgGs induces severe lung injury by skewing macrophage polarization away from wound-healing ‘M2’ characteristics towards a strong pro-inflammatory phenotype. Notably, SARS patients that eventually died from infection displayed similar conversion of these wound-healing lung macrophages, as well as the early and high presence of neutralizing IgG antibodies. The pro-inflammatory consequences of these IgGs could be suppressed by Fc receptor (FcR) blockade. Very similar to SARS, severe COVID-19 patients are characterized by an early rise and high titers of IgG antibodies(*10–12*) and show a similar conversion from anti- to pro-inflammatory lung macrophages(*13*). Combined, these data hint towards the involvement of anti-Spike IgG in severe cases of COVID-19. Therefore, in this study we explored the hypothesis that, similar to responses in SARS, anti-Spike antibodies drive excessive inflammation in severe cases of COVID-19.

We assessed the effect of anti-Spike antibodies from serum of critically ill COVID-19 patients on human M2 macrophages. While different protocols are available for generating human M2 macrophages, our previous transcriptional analyses demonstrated that an M-CSF and IL-10-induced monocyte differentiation protocol generates cells that most closely resemble primary human lung macrophages(*14*). Since activation of immune cells by IgG antibodies is known to require immune complex formation by binding of IgG to its ligand(*15, 16*), we generated Spike-IgG immune complexes by incubating SARS-CoV-2 Spike-coated wells with diluted serum from severely ill COVID-19 patients (i.e. patients from the intensive care unit at the Amsterdam UMC) that tested positive for anti-SARS-CoV-2 IgG. As shown in Figure 1A, stimulation with Spike protein alone did not induce cytokine production, while Spike-IgG immune complexes elicited small amounts IL-1β, IL-6, and TNF, but very high IL-8 production by human macrophages. Stimulation with the viral mimic PolyIC induced very little cytokine secretion. Yet, since in the late phase of infection lung macrophages are simultaneously exposed to viral stimuli and anti-Spike IgG immune complexes, we also assessed the effect of the combination of these two stimuli. Strikingly, combined stimulation of anti-Spike IgG immune complexes and PolyIC strongly amplified the production of COVID-19-associated pro-inflammatory cytokines IL-1β, IL-6, and TNF (Figure 1A). Induction of the anti-inflammatory cytokine IL-10 was also increased, similar to what is observed in COVID-19 patients(*17*). To verify the relevance of our findings for human lung macrophages, we assessed the response of primary alveolar macrophages that were obtained by bronchoalveolar lavage, which showed very similar responses (Figure 1B).

**Fig. 1.**
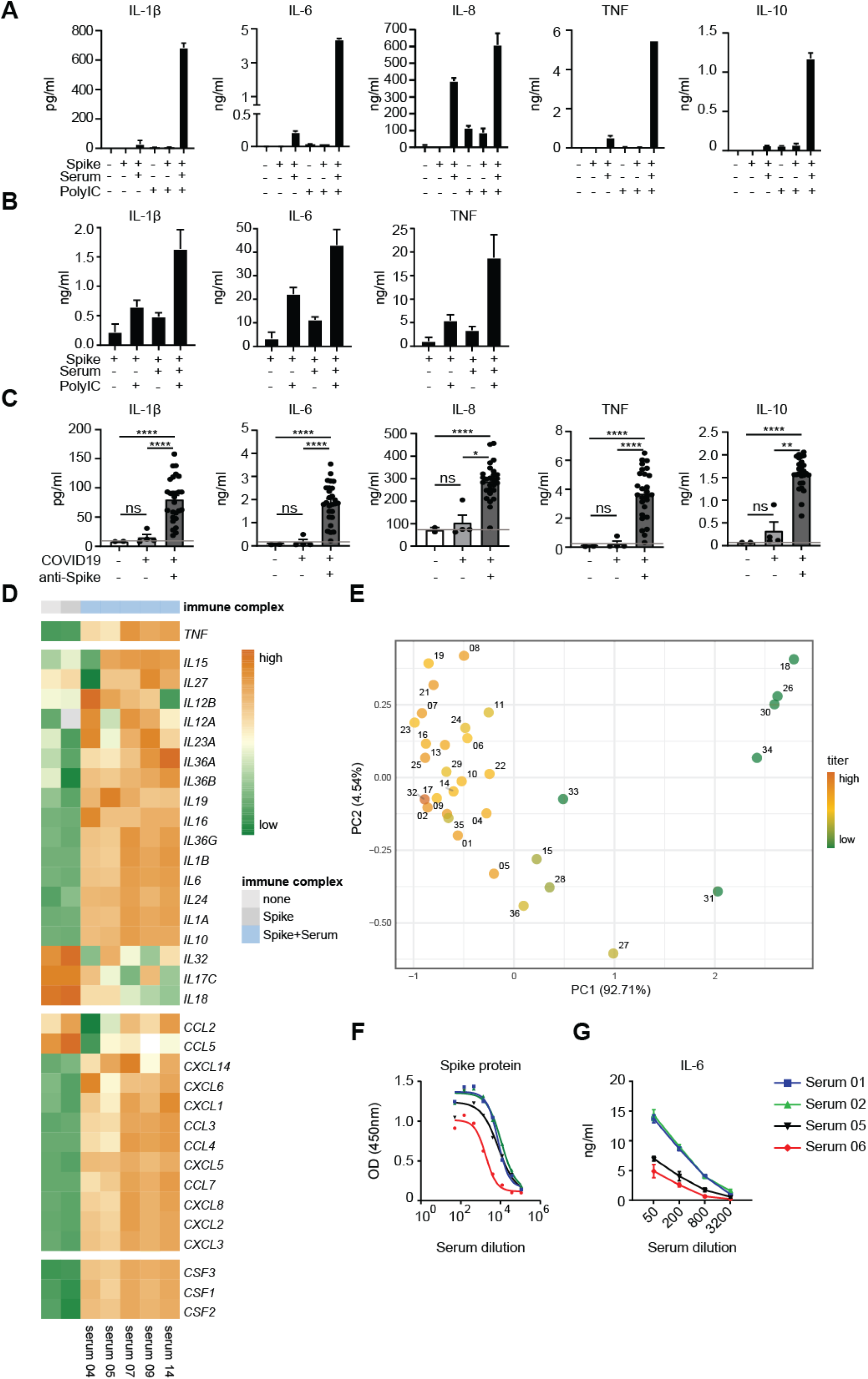
Anti-Spike IgG induces inflammation by macrophages. (A) Cytokine production by human macrophages after 24h stimulation with (combinations of) Spike protein, COVID-19 serum (50x diluted), and PolyIC. Triplicate values from a representative experiment with serum from five different COVID-19 patients and two different macrophage donors (mean+stdev). (B) Cytokine production by primary alveolar macrophages obtained from a bronchoalveolar lavage, as stimulated under A. Triplicate values from an experiment with one donor (mean+stdev). (C) Macrophages stimulated with Spike protein and PolyIC were co-stimulated with serum from ICU lung patients that either did not have COVID-19, had COVID-19 but were negative for anti-Spike IgG, or had COVID-19 and were positive for anti-Spike IgG. Every dot represents cytokine production after 24h by a different serum donor (mean+SEM). Horizontal grey line depicts cytokine induction upon stimulation with PolyIC+Spike protein. *P<0.05; **P<0.01; ****P<0.0001; ns=not significant. (D) Heatmap showing scaled log_2_ expression level of genes assessed by RNAseq upon 6h stimulation of human macrophages with PolyIC, with or without Spike protein and serum from five COVID-19 patients that tested positive for anti-Spike IgG. (E) Principle component analysis of the combined cytokine profile (IL-1β, IL-6, IL-8, IL-10, TNF, IFN-β, IFN-γ, CXCL10) for all serum samples overlaid with log_10_ IgG titers. Numbers represent the serum number (see Table S1 for details). (F) Binding ELISA to determine the relative titer of anti-Spike IgG in sera of four severe COVID-19 patients. OD=optical density. (G) Macrophages stimulated with Spike protein and PolyIC were co-stimulated with different dilutions of serum from patients in panel F, after which IL-6 production was determined after 24h.

To assess whether this inflammatory response is really dependent on anti-Spike IgG antibodies, and not on other inflammatory components in serum of critically ill patients, we compared the effect of sera from 33 intensive care lung disease patients that either (1) did not have COVID-19, (2) had COVID-19 but were still negative for anti-Spike IgG, or (3) had COVID-19 and were positive for anti-Spike IgG (see Table S1 for details). While serum of non-COVID-19 patients and anti-Spike IgG-negative COVID-19 patients showed no up-regulation of pro-inflammatory cytokines compared to individual PolyIC stimulation, IL-1β, IL-6, IL-8, and TNF production was strongly amplified by serum of COVID-19 patients with anti-Spike IgG (Figure 1C). Substantiating these findings, RNAseq analysis of macrophages stimulated with sera from anti-Spike IgG positive COVID-19 patients also showed clear induction of a pro-inflammatory gene program, as highlighted by induction of *TNF*, interleukins, chemokines, and macrophage differentiation factors (Figure 1D). Interestingly, also IFN-β and IFN-γ were induced by anti-Spike positive serum, while the classical downstream interferon response gene CXCL10 was reduced (Figure S1A and S1B), which may be related to reduced expression of IFN receptors (Figure S1C).

In COVID-19 patients, high anti-Spike IgG titers are associated with disease severity(*10, 12*). To determine whether anti-Spike titers correlate with higher cytokine responses by human macrophages, we performed a principle component analysis (PCA) of the combined cytokine production data for all samples, which upon overlaying with anti-Spike IgG titers (based on half maximal effective concentration EC_50_) indeed indicated that the hyper-inflammatory response of macrophages was associated with IgG titers (Figure 1E). This was subsequently confirmed by analyzing four serum samples with different titers using serial-step dilutions (Figure 1F), which showed dose-dependent induction of pro-inflammatory cytokines (Figure 1G). Taken together, these data demonstrate that anti-Spike IgG immune complexes generated from serum of severely ill COVID-19 patients induce a strong pro-inflammatory response by (otherwise immunosuppressive) human M2 macrophages, which is characterized by high production of classical cytokine storm mediators such as IL-1β, IL-6, IL-8, and TNF.

In addition to excessive lung inflammation through a cytokine storm, severe COVID-19 is characterized by pulmonary edema, following disruption of the microvascular endothelium(*18*), and by coagulopathy, which in many patients is characterized by pulmonary thrombosis(*19*). To test whether the excessive macrophage activation by anti-Spike IgG may contribute to pulmonary edema and thrombosis, we applied in vitro models for endothelial barrier integrity(*20*) and in situ thrombosis(*21*) using primary human pulmonary artery endothelial cells (HPAEC), where thrombocytes can be added under flow conditions. While conditioned medium of macrophages that had been stimulated with only PolyIC induced a transient drop in endothelial barrier integrity, co-stimulation of macrophages with Spike protein and serum of severe COVID-19 patients induced long-lasting endothelial barrier disruption (Figure 2A). In addition, during platelet perfusion we observed significantly increased platelet adhesion to endothelium exposed to conditioned medium of macrophages that had been co-stimulated with Spike protein and serum (Figure 2B). This effect was paralleled by an increase in von Willebrand Factor release from the endothelial cells (Figure 2C), indicative for an active pro-coagulant state of the endothelium. These data indicate that anti-Spike IgG from severely ill COVID-19 patients does not only induce hyperinflammation by macrophages, but also may contribute to permeabilization of pulmonary endothelium and microvascular thrombosis.

**Fig. 2.**
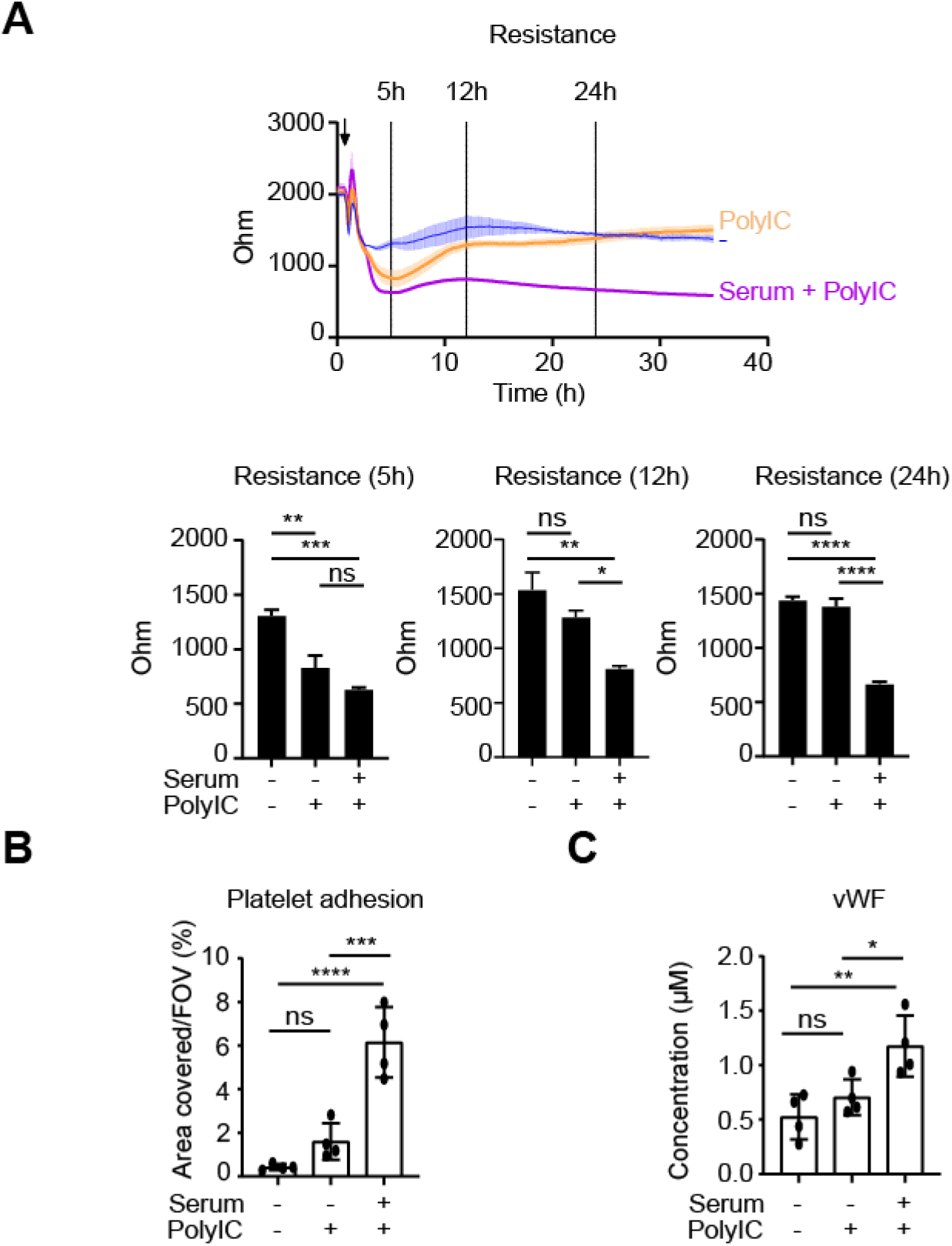
Anti-Spike IgG breaks endothelial barrier integrity and activates platelets. (A) Human pulmonary arterial endothelial cells were exposed to supernatant of macrophages that were unstimulated, or had been stimulated with PolyIC and Spike protein, with or without COVID-19 serum. Endothelial barrier integrity was measured over time by measuring the resistance over time using electrical cell-substrate impedance sensing. (B) Endothelium as stimulated under A. for 24h was perfused with platelets for 5 minutes, after which the area covered by platelets was quantified. (C) Flow supernatant was collected after perfusion under B, and VWF levels were measured with ELISA. *P<0.05; **P<0.01; ***P<0.001; ****P<0.0001; ns=not significant.

In addition to the anti-Spike antibodies from serum, we tested the effect of the recombinant anti-Spike IgG COVA1-18, which we generated previously from B cells of COVID-19 patients(*22*). As previously shown by others, the concentration of anti-SARS-CoV-2 IgG in severe COVID-19 patients peaks at an average of 16.5 μg/mL(*23*). To test the efficacy of our recombinant anti-Spike IgG, we stimulated macrophages with anti-Spike immune complexes made with a high concentration (mimicking a serum concentration of 100 μg/mL in our assay) of the recombinant antibody COVA1-18. Remarkably, the high concentration of recombinant anti-Spike IgG immune complexes elicited substantially less IL-1β, IL-6, and TNF than anti-Spike immune complexes made from COVID-19 serum (Figure 3A). Interestingly, we did not observe this difference for the induction of anti-inflammatory cytokine IL-10 (Figure 3A). These data suggest that the anti-Spike IgG in severe cases of COVID-19 patients is intrinsically more pro-inflammatory than recombinant IgG. One of the key characteristics that determines IgG pathogenicity is glycosylation of the IgG Fc tail at position 297(*24, 25*). Previously, we and others have shown that anti-Spike IgG of severe COVID-19 patients has aberrant fucose and galactose expression, both compared to the total IgG within these individual patients, as well as compared to anti-Spike IgG from mild or asymptomatic patients(*26, 27*). For a subset of COVID-19 serum samples in the present study, the glycosylation pattern of anti-Spike IgG1 had been determined previously for the study of Larsen et al.(*26*), which showed significantly decreased fucosylation and increased galactosylation of anti-Spike IgG compared to total IgG within the tested patients (IgG glycosylation of the sera used in this study is depicted in Figure S2A). When plotting the percentage of anti-Spike IgG fucosylation against pro-inflammatory cytokine production, we observed an inverse correlation for IgG fucosylation and production of IL-1β, IL-6, and TNF (Figure 3B), while this was not seen for IL-8 and IL-10 (Figure S2B). To determine whether there is a causative role for these glycosylation changes in the induction of pro-inflammatory cytokines, we compared cytokine induction by regular COVA1-18, to modified COVA1-18 that had low fucose and high galactose (glycosylation details in Table S2). Notably, COVA1-18 with low fucose and high galactose showed an increased capacity for amplification of pro-inflammatory cytokines (Figure 3C). These data indicate that anti-Spike IgG from COVID-19 patients has an aberrant glycosylation pattern that makes these antibodies intrinsically more inflammatory than ‘common’ IgGs by increasing its capacity to induce high amounts of pro-inflammatory cytokines.

**Fig. 3.**
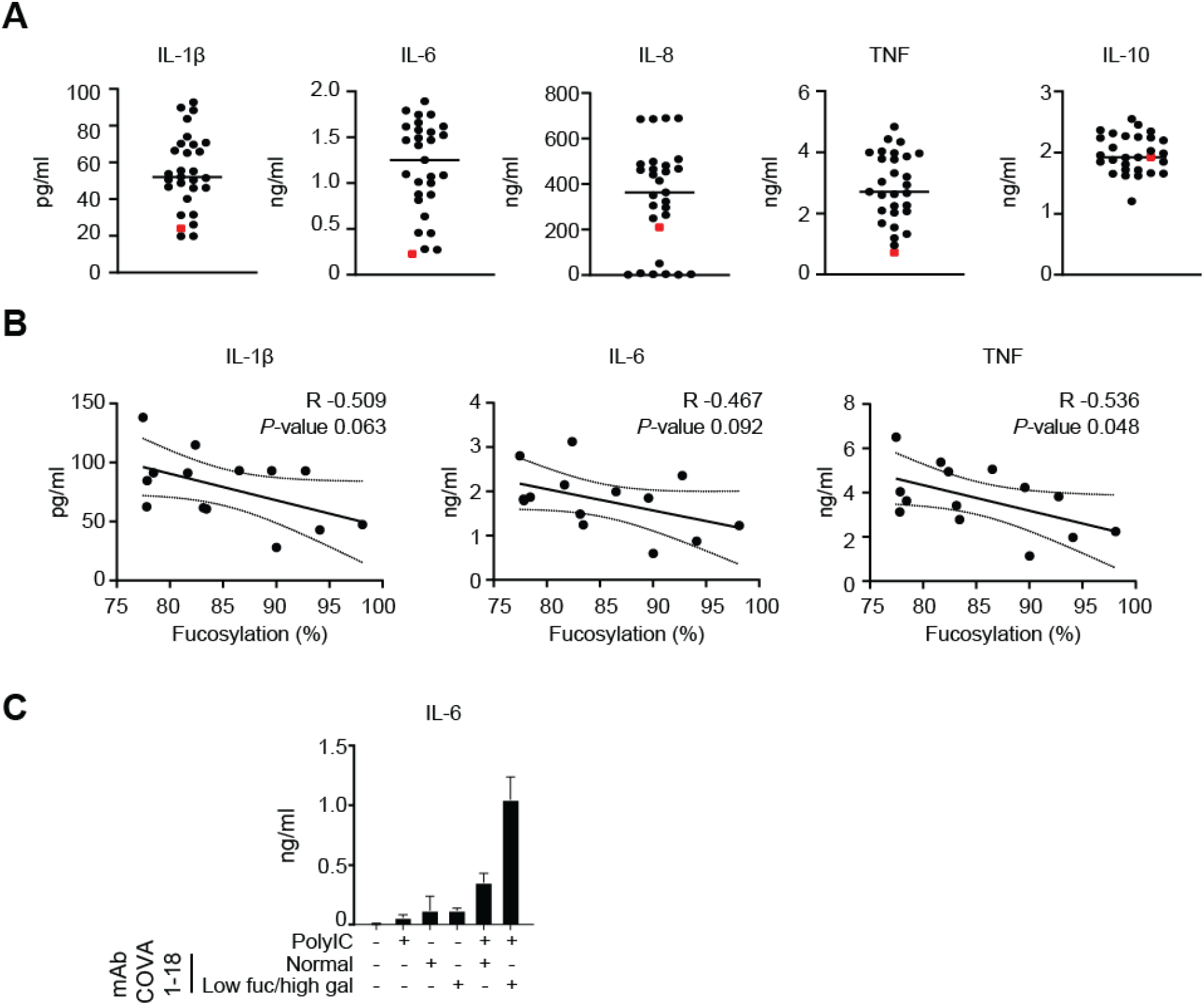
Anti-Spike IgG-induced inflammation is dependent on IgG Fc glycosylation. (A) Macrophages stimulated with Spike protein and PolyIC were co-stimulated with either 50x diluted serum from different anti-Spike positive COVID-19 patients (black dots), or with recombinant anti-Spike antibody COVA1-18 (red dot). Cytokine production was measured after 24h. (B) Correlation graphs of fucosylation percentages of anti-Spike IgG1 from COVID-19 serum against cytokine production of macrophages after stimulation as in panel A. The Pearson correlation coefficient (R) and *P* value are illustrated on each graph with 95% confidence bands of the best-fit line. (C). Macrophages stimulated with Spike protein were co-stimulated with (combinations of) PolyIC, COVA1-18 (recombinant anti-Spike IgG1), or COVA1-18 that had been modified to express low fucose and high galactose. IL-6 production was measured after 24h. Representative donor (mean+stdev) of 4 independent experiments.

Anti-Spike IgG from severely ill COVID-19 patients promoted inflammatory cytokines, endothelial barrier disruption, and microvascular thrombosis, which are key phenomenon in severely ill COVID-19 patients that are thought to underlie the pathology. Hence, counteracting this antibody-induced aberrant immune response could be of potential therapeutic interest. To determine how this antibody-induced inflammation could be counteracted, we first set out to investigate which receptors on human macrophages are activated by the anti-SARS-CoV-2 IgG immune complexes. IgG immune complexes can be recognized by Fc gamma receptors (FcγRs), which includes FcγRI, FcγRII, and FcγRIII(*16*). As shown in Figure 4A, the used human macrophage model highly expressed all FcγRs. To determine whether FcγRs are involved in activation by anti-Spike immune complexes, we blocked the different FcγRs with specific antibodies during stimulation, and analyzed cytokine production. As shown in Figure 4B, all FcγRs contributed to anti-Spike-induced cytokine induction, but the most pronounced inhibition was observed upon blockade of FcγRII. No inhibition was observed upon blocking of Fc alpha receptor I (FcαRI), suggesting that IgA does not play a major role in the observed cytokine induction (Figure 4B).

**Fig. 4.**
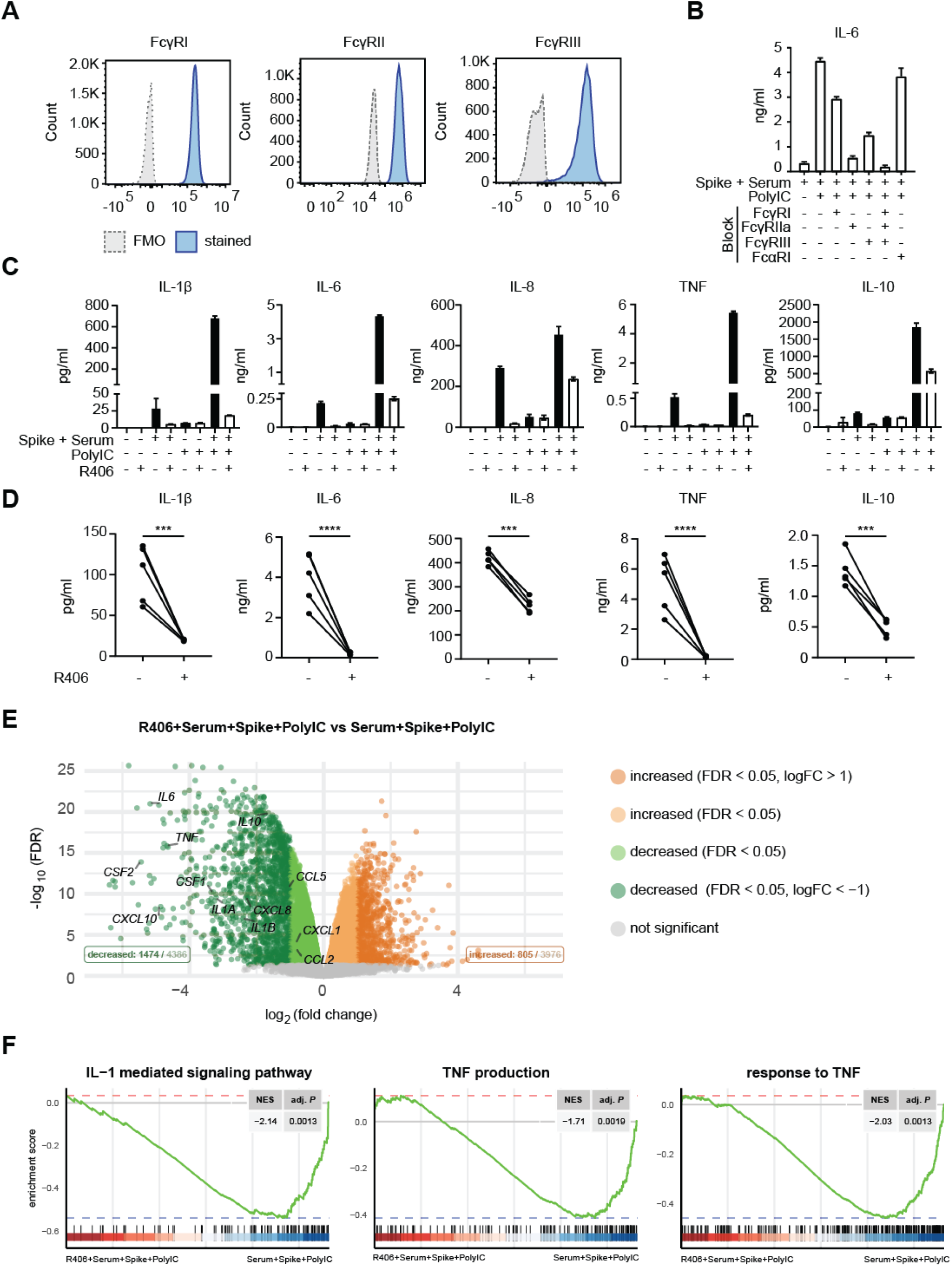
Anti-Spike IgG-induced inflammation is counteracted by fostamatinib. (A) Membrane expression of FcγRI, II, and III by human macrophages was determined by flow cytometry. (B) FcγRI, II, and/or III, or FcαRI were blocked by specific antibodies, after which macrophages were stimulated with Spike, COVID-19 serum, PolyIC, or a combination. IL-6 production was measured after 24h. Triplicate values from a representative experiment with serum from three different COVID-19 patients and two different macrophage donors (mean+stdev). (C/D). Macrophages were pre-incubated with Syk inhibitor R406, after which cells were stimulated as in B. Cytokine production was measured after 24h. Panel C shows a representative donor (mean+stdev), panel D shows the response of macrophages stimulated with Spike, PolyIC, and COVID-19 serum, with or without pre-incubation with R406. Every pair of dots represents cytokine production after 24h by a different serum donor. ***P<0.001; ****P<0.0001. (E) Volcano plot depicting up- and down-regulated genes when comparing macrophages stimulated for 6h with Spike, PolyIC, and serum to the same stimulation in the presence of R406. FDR=false discovery rate. (F) Gene set enrichment analysis (GSEA) of curated gene sets suppressed by R406: interleukin-1-mediated signaling pathway (GO:0070498), TNF production (GO:0032640), response to TNF (GO:0034612). NES stands for normalized enrichment score and adj. *P* represents the BH-adjusted *P* value.

FcγRs are known to induce signaling that critically depends on the kinase Syk(*28, 29*). To determine whether we could counteract anti-Spike-induced immune activation, we blocked Syk using R406, the active component of the small molecule inhibitor fostamatinib, an FDA- and EMA-approved drug for treatment of immune thrombocytopenia (ITP)(*30*). Strikingly, R406 almost completely blocked pro-inflammatory cytokine production induced by anti-Spike IgG from severe COVID-19 patients (Figure 4C and D). Importantly, inhibition by R406 appeared to be specific, since it selectively blocked anti-Spike induced amplification of cytokines, but did not substantially affect cytokine production induced by PolyIC alone (Figure 4C).

To assess the effects of inhibition by fostamatinib in greater detail, we analyzed the effects of R406 on macrophages stimulated with Spike, COVID-19 serum, and PolyIC by RNAseq. In total 4386 genes were suppressed by R406 treatment, while 3976 genes were induced (FDR<0.05, Figure 4E). In the downregulated genes many of the classical pro-inflammatory mediators were present, including *TNF*, *IL1B*, *IL6* and *CCL2*. Pathway analyses further showed no clear pathways in the upregulated genes, while suppressed genes were clearly linked to inflammatory pathways (Figure S3A). Finally, gene set enrichment analysis (GSEA) showed that genes associated with several pro-inflammatory pathways including IL-1 signaling and TNF production and response, were significantly downregulated by R406 (Figure 4F), while also response to type I IFN, Fc-gamma receptor signaling, glycolysis, and platelet activation gene sets were suppressed (Figure S3B). These data demonstrate that the excessive inflammatory response by anti-Spike IgG from severely ill COVID-19 patients can be counteracted by the Syk inhibitor fostamatinib.

In conclusion, our data show that anti-Spike IgG from serum of severely ill COVID-19 patients strongly amplifies pro-inflammatory responses by human macrophages, and can contribute to subsequent endothelial barrier disruption and thrombosis. This may explain the observation that many COVID-19 patients become critically ill around the time of activation of adaptive immune responses. In general, antibodies are beneficial for host defense by providing various mechanisms to counteract infections(*31*). These different effector functions of antibodies are modulated by antibody-intrinsic characteristics, such as isotype, subclass, allotype, and glycosylation(*24*). In severely ill COVID-19 patients the glycosylation of anti-Spike IgG is changed, which we here show amplifies the IgG effector function to promote pro-inflammatory cytokine production, of which COVID-19-associated cytokines such as IL-1β, IL-6, and TNF(*5, 32*) are most pronounced. Decreased IgG fucosylation, as observed in severe cases of COVID-19, has so far only been observed in patients infected with HIV and Dengue virus(*33, 34*), but may actually be a general phenomenon in a response to enveloped viruses(*26*). While decreased fucosylation increases the infection of cells by a process known as antibody-dependent enhancement (ADE)(*35*), there is little evidence for antibody-enhanced infection in COVID-19(*36*). Instead, our data show that increased pathology by de-fucosylated IgG in COVID-19 patients most likely results from excessive immune activation. The combination of decreased fucosylation and increased galactosylation of IgG is known to particularly increase the affinity for FcγRIII(*24*). While we show that FcγRIII was partially responsible for anti-Spike-induced inflammation, FcγRII contributed most, indicating that collaboration between multiple FcγRs is required for the hyper-inflammatory responses induced by the aberrant glycosylation of anti-Spike IgG. In addition to human alveolar macrophages, these FcγRs are expressed by various other myeloid immune cells(*15*), but also by airway epithelial cells(*37*), which are one of the main target cells of infection by SARS-CoV-2 and closely interact with activated macrophages(*38*).

The observed hyper-inflammatory response induced by anti-Spike IgG from severe patients could be specifically counteracted by the Syk inhibitor R406, the pro-drug of fostamatinib. Notably, fostamatinib is an FDA and recently also EMA approved drug that is currently used for treatment of ITP(*30*), which may facilitate repurposing for the treatment of severe COVID-19 patients. A very recent study indicates that fostamatinib may also counteract acute lung injury by inhibiting Mucin-1 expression on epithelial cells, suggesting that fostamatinib may target multiple pathways simultaneously(*39*). In addition to fostamatinib, also other drugs that target key molecules in FcγR signaling could be efficacious to counteract anti-Spike IgG-induced inflammation in COVID-19 patients. For example, the Syk-dependent FcγR signaling pathway critically depends on the transcription factor IRF5(*15, 28*), which can be targeted using cell penetrating peptides(*40*). Furthermore, FcγR stimulation is known to induce metabolic reprogramming of human macrophages(*28*), which is also observed in COVID-19 patients(*41*), and therefore may provide additional targets for therapy. These findings may not only be valuable to find new ways to treat the most severely ill COVID-19 patients, but may also have implications for the therapeutic use of convalescent serum, for which it may be wise to omit the de-glycosylated IgGs that are present in severely ill patients.

## Author contributions

Conceptualization: JdD, MdW, GV

Methodology: WH, HJC, SA, XM, JA, HJB, MvG, GV, MdW, JdD, MW, FL, SdT, ML, PB, TC, KvS, MvG

Formal analysis: HJC, WH, SA, XM, JA, MW, MvG, ML

Investigation: WH, HJC, SA, XM, SdT, KG, GG, AN, LW

Resources: PB, TC, KvS, SdT, GdB, CvD, MvG, GV, PB, FvB, RJ, AV, Amsterdam UMC COVID-19 Biobank

Data Curation: HJC, MdW, MW, GV, ML

Writing – original draft: JdD

Writing – review and editing: MdW, WH, HJC, SA, XM, JA, PB, SdT, AV, RS, HJB, MvG, GV

Visualization: HJC, MdW, WH, SA, GV

Supervision: JdD, MdW, MvG, GV, JA, HJB, CvD, AV, GdB, RS, MW, TC

Funding acquisition: JdD; MdW; MvG; GV

## Data and materials availability

We will share reagents, materials and data presented in this study on request. The raw RNASeq data will be stored in the European Genome-Phenome Archive. The accession number and associated data will be made available after publication to all researchers upon request to the Data Access Committee. The restriction on data access is required for human donor protection.

## Supplementary Materials

### Materials and Methods

#### Cells

Buffy coats from healthy anonymous donors were acquired from the Sanquin blood supply in Amsterdam, the Netherlands. All the subjects provided written informed consent prior to donation to Sanquin. Monocytes were isolated from the Buffy coats through density centrifugation using Lymphoprep™ (Axis-Shield) followed by human CD14 magnetic beads purification with the MACS^®^ cell separation columns (Miltenyi) as previously described(*14*). The resulting monocytes were seeded on tissue culture plates and subsequently differentiated to macrophages for 6 days in the presence of 50ng/mL human M-CSF (Miltenyi) with Iscove’s Modified Dulbecco’s Medium (IMDM, Lonza) containing 5% fetal bovine serum (FBS) (Biowest) and 86μg/mL gentamicin (Gibco). The medium was renewed on the third day. After 6-day differentiation, the medium was replaced by culture medium without M-CSF and supplemented with 50 ng/mL IL10 (R&D) for 24 hours to generate alveolar macrophage-like monocyte-derived macrophages. These macrophages were then detached with TrypLE Select (Gibco) for further treatment and stimulation.

Pulmonary artery endothelial cells (PAEC) were obtained from resected pulmonary artery tissue, obtained from lobectomy surgery performed at Amsterdam UMC, location VU University Medical Center, The Netherlands and isolated according to the previously published protocol(*21*). Briefly, the endothelial cell layer was carefully scraped onto fibronectin-coated (5 μg/mL) culture dishes (Corning, #3295), and maintained in culture in endothelial cell medium (ECM, ScienCell, #1001) supplemented with 1% Penicillin/Streptomycin, 1% endothelial cell growth supplement (ECGS), 5% FBS, and 1% non-essential amino acids (NEAA, Biowest, #X055-100). Cells were grown until passage 4-6 for experiments.

Primary macrophages were prepared from broncheo-alveolar lavage (BAL) fluid that was obtained as spare material from the ongoing DIVA study (Netherlands Trial Register: NL6318; AMC Medical Ethical Committee approval number: 2014_294). The DIVA study includes healthy male volunteers aged 18-35. In this study, the subjects are given a first hit of lipopolysaccharide (LPS) and, two hours later, a second hit of either fresh or aged platelet concentrate or NaCl 0.9%. Six hours after the second hit, a BAL is performed by a trained pulmonologist according to national guidelines. Fractions 2-8 are pooled and split in two, one half is centrifuged (4 °C, 1750g, 10 minutes), the cell pellet of which was used in this research. Since the COVID-19 pandemic, subjects are also screened for SARS-CoV-2 (via throat swab PCR) 2 days prior to the BAL. All subjects in the DIVA study have signed an informed consent form. The amount of macrophages (80-85%) in the BAL was determined by counting the cells that did not adhere to the plate after 30 minutes at 37°C. For our experiment complete cell pellet was stimulated.

#### Coating

To mimic Spike protein specific immune complexes, 2μg/mL soluble prefusion-stabilized Spike proteins of SARS-CoV-2 was coated overnight on a 96-well high-affinity plate (Nunc). Plate were blocked with 10% FCS in PBS for 1h at 37°C. Then diluted serum or 2 g/mL anti-SARS-CoV-2 monoclonal antibodies were added and incubated for 1h at 37°C. The Spike and anti-SARS-CoV-2 monoclonal antibody COVA1-18 were generated as described previously(*22*). Patient sera were provided by the Amsterdam UMC COVID-19 Biobank based on a deferred consent procedure for the usage of materials and clinical information for research purposes, approved by the medical ethics committees of Amsterdam University Medical Centers (Amsterdam, The Netherlands).

The specific glyco-engineered antibodies were made from the potent SARS-CoV-2 neutralizing antibody COVA1-18 produced in 293F cells as previously described(*22*). Glyco-engineering tools were used to alter N-linked glycosylation of the N297 glycan in the Fc domain and thereby generated several COVA1-18 glycoforms(*42*). To decrease fucosylation of the N-linked glycan 0.2mM of the decoy substrate for fucosylation, 2-deoxy-2-fluoro-l-fucose (2FF) (Carbosynth, MD06089) was added one hour prior to transfection. To produce a COVA1-18 variant with elevated galactosylation, 293F cells were co-transfected (1% of total DNA) with a plasmid expressing Beta-1,4-Galactosyltransferase 1 (B4GALT1). In addition 5mM D-Galactose was added 1h before transfection. Antibodies were purified with protein G affinity chromatography as previously described(*22*) and stored in PBS at 4°C. To determine the glycosylation of COVA1-18, aliquots of the mAb samples (5μg) were subjected to acid denaturation (100 mM formic acid, 5 minutes), followed by vacuum centrifugation. Subsequently, samples were trypsinized, and Fc glycopeptides were measured as described previously(*26*). Relative abundances of Fc glycopeptides were determined, and levels of bisection, fucosylation, galactosylation and sialylation were determined as described before(*26*).

#### Stimulation

Cells (50,000/well) were stimulated in a pre-coated plates as described above in combination with 20 μg/mL PolyIC (Sigma-Aldrich). To block Syk, cells were pre-incubated with 0.5 μM R406 (Selleckchem) or DMSO as a control, for 30 minutes at 37°C. To block the different FcRs, cells were pre-incubated with 20 μg/mL of the following antibodies: (anti-FcyRI (CD64; 10.1; BD Bioscience); anti-FcyRIIa (CD32a; IV.3; Stemcell Technologies); anti-FcyRIII (CD16; 3G8; BD Bioscience) and anti-FcαRI (CD89; MIP8a; Abcam)) for 30 minutes at 4°C. Then media was added to get a final antibody concentration of 5 μg/mL.

#### Endothelial barrier function

PAEC passage 4-6 were seeded 1:1 in 0.1% gelatin coated 8-well (8W10E) or 96-well (96W10idf PET) IBIDI culture slides for electrical cell-substrate impedance sensing (ECIS), as previously described(*20*). Cells were maintained in culture in endothelial cell medium (ECM, ScienCell, #1001) supplemented with 1% Penicillin/Streptomycin, 1% endothelial cell growth supplement (ECGS), 5% FBS and 1% non-essential amino acids (NEAA, Biowest, #X055-100), with medium change every other day. From seeding onwards, electrical impedance was measured at 4000Hz every 5 minutes. Cells were grown to confluence and after 72h, ECM medium was removed and replaced by supernatant of alveolar macrophage-like monocyte-derived macrophages stimulated for 6h as described above with PolyIC, or in combination with patient serum. Within every experiment triplicate measurements were performed for each condition. For every experiment PAECs and macrophages obtained from different donors were used.

#### Platelet adhesion on PAEC under flow

PAECs were seeded in 0.1% gelatin coated μ-Slide VI 0.4 ibiTreat flow slides (ibidi, #80606) and cultured for 7 days. PAECs were pre-incubated for 24h with supernatant of alveolar macrophage-like monocyte-derived macrophages stimulated for 6h as described above with PolyIC, or in combination with patients serum before flow experiments were performed. On the day of perfusion, citrated blood was collected from healthy volunteers and platelets were isolated as previously described(*43*). Platelets were perfused for 5 minutes and phase-contrast and fluorescent images were taken with an Etaluma LS720 microscope using a 20X phase-contrast objective. Platelet adhesion was quantified in ImageJ by determining the area covered by platelets per Field of View (FOV).

#### ELISA

To determine cytokine production, supernatants were harvested after 24h of stimulation and cytokines were detected using the following antibody pairs: IL-1β and IL-6 (U-CyTech Biosciences); TNF (eBioscience); and IL-8 (Invitrogen).

To determine the concentration of anti-Spike antibodies present in patients serum, Spike protein was coating directly on 96-well plates at 5μg/mL overnight to determine serum binding using serial dilutions as previously described(*22*).

To detect the VWF levels, flow supernatant was collected after perfusion and VWF levels were measured with ELISA. An 96-well high affinity ELISA plate was coated with polyclonal anti-VWF (1:1000, Dako, #A0082) and blocked with 2% BSA. Samples were loaded and bound VWF was detected with HRP-conjugated rabbit polyclonal anti-VWF (1:2500, Dako, #A0082). Normal plasma with a stock concentration of 50 nM VWF (gifted from Sanquin) was used as a standard for determination of VWF levels, measured at 450nm and 540nm.

#### Meso Scale Discovery multiplex assay

V-PLEX Custom Human Cytokine10-plex kits for Proinflammatory Panel1 and Chemokine Panel 1 (K151A0H-2, for IL-1β, IL-6, IL-8, IL-10, TNF, CCL2, CXCL10) and U-PLEX human Interferon Combo SECTOR (K15094K-2, for IFN-α2a, IFN-β, IFN-γ, IFN-λ1) were purchased from Mesoscale Discovery (MSD). The lyophilized cocktail mix calibrators for Proinflammatory Panel 1, Chemokine Panel 1, and 4 calibrators for U-PLEX Biomarker Group 1 (Calibrator 1, 3, 6, 9) were reconstituted in provided assay diluents respectively. U-PLEX plates were coating with supplied linkers and biotinylated capture antibodies according to manufacturer’s instructions. Proinflammatory cytokines and chemokines in supernatant collected at 24 hours after stimulation were detected with pre-coated V-PLEX while interferons in 6-hour supernatant were measured by coated U-PLEX plates. The assays were performed according to manufacturer’s protocol with overnight incubation of the diluted samples and standards at 4°C. The electrochemiluminescence signal (ECL) were detected by MESO QuickPlex SQ 120 plate reader (MSD) and analyzed with Discovery Workbench Software (v4.0, MSD). The concentration of each sample was calculated based on the four-parameter logistic fitting model generated with the standards (concentration was determined according to the certificate of analysis provided by MSD). log_10_ values of measured level of IL-1β, IL-6, IL-8, IL-10, TNF, IFN-β, IFN-γ, CXCL10 were used for the principle component analysis. log_10_ IgG titers (half maximal effective concentration, EC_50_) were used for the color overlay.

#### RNA Sequencing

Cells were stimulated as described above and lysed after 6 hours. Total RNA was isolated with RNeasy Mini Kit (Qiagen) and RNase-Free DNase Set (Qiagen) per the manufacturer’s protocol. cDNA libraries were prepared using the standard protocol of KAPA mRNA HyperPrep Kits (Roche) with input of 300ng RNA per sample. Size-selected cDNA libraries were pooled and sequenced on a HiSeq 4000 sequencer (Illumina) to a depth of 16-20M per sample according to the 50 bp single-end protocol at the Amsterdam University Medical Centers, location Vrije Universiteit medical center. Raw FASTQ files were aligned to the human genome GRCh38 by STAR (v2.5.2b) with default settings(*44*). Indexed Binary alignment map (BAM) files were generated and filtered on MAPQ>15 with SAMTools (v1.3.1)(*45*). Raw tag counts and reads per kilo base million (RPKM) per gene were calculated using HOMER2’s analyzeRepeats.pl script with default settings and the -noadj or -rpkm options for raw counts and RPKM reporting(*46*) for further analyses.

#### Flow cytometry

After detachment, macrophages were stained with antibodies against Fc gamma receptors: FcγRI (CD64; cat# 305014, Biolegend), FcγRII(CD32; cat# 555448, BD bioscience), and FcγRIII (CD16; cat# 562293, BD bioscience). Fluorescence was measured with CytoFLEX Flow Cytometer and analyzed with FlowJo software version 7.6.5 (FlowJo, LLC, Ashland, OR). Fluorescence Minus One (FMO) controls were used for each staining as negative controls.

#### Data availability

All data generated during this study are available within the paper. The RNAseq data were deposited on Gene Expression Omnibus. Source data are provided with this paper.

#### Statistical analysis

Statistical significance of the data was performed in Graphpad Prism version 8 (GraphPad Software). Correlation analysis: for the fucosylation and cytokine correlation analysis, Pearson’s correlation was performed for paired fucosylation percentage and measured cytokine concentration. For Fig. 1C and Fig. S1B, statistics were made with Brown-Forsythe and Welch’s ANOVA test and corrected by Dunnett T3 test for multiple test correction (sample size: COVID19−: 2, COVID19+ anti-spike−: 4, COVID19+ anti− spike+: 27). For Fig. 2 an ordinary one-way ANOVA was performed and corrected for Tukey’s comparisons test (Fig 2A) or Sidak’s multiple comparison test (Fig. 2B and C). For Fig. 4D and Fig. S2A ratio paired t-test were used.

#### Functional analyses of transcriptomic data

All analyses were performed in the R statistical environment (v3.6.3). Differential expression was assessed using the Bioconductor package edgeR (v3.28.1)(*47*). Lowly expressed genes were filtered with the filterByExpr function and gene expression called differential with a false discovery rate (FDR) <0.05. Pathway enrichment analyses were performed on the differentially regulated genes with an absolute log_2_(fold change) higher than 1 using the Metascape (http://metascape.org/gp/index.htmL) (*48*) on 2020-06-26. For heatmaps, normalized expression values (count per million, CPM) of each gene were calculated and plotted using pheatmap (v1.0.12) with values scaled by gene. Gene set enrichment analysis (GSEA) was performed with Bioconductor package fgsea (v1.12.0)(*49*) with genes ranked by effect size (Cohen’s *d*) with respect to the “R406+serum+spike+PolyIC vs serum+spike+PolyIC” against the curated gene sets obtained from gene ontology (GO) by Bioconductor package biomaRt (v2.42.1)(*50*). A total of 5000 permutations were performed to estimate the empirical *P* values for the gene sets. Normalized enrichment scores and the BH-adjusted *P* values were provided in the figure.

---

**Figure S1.**
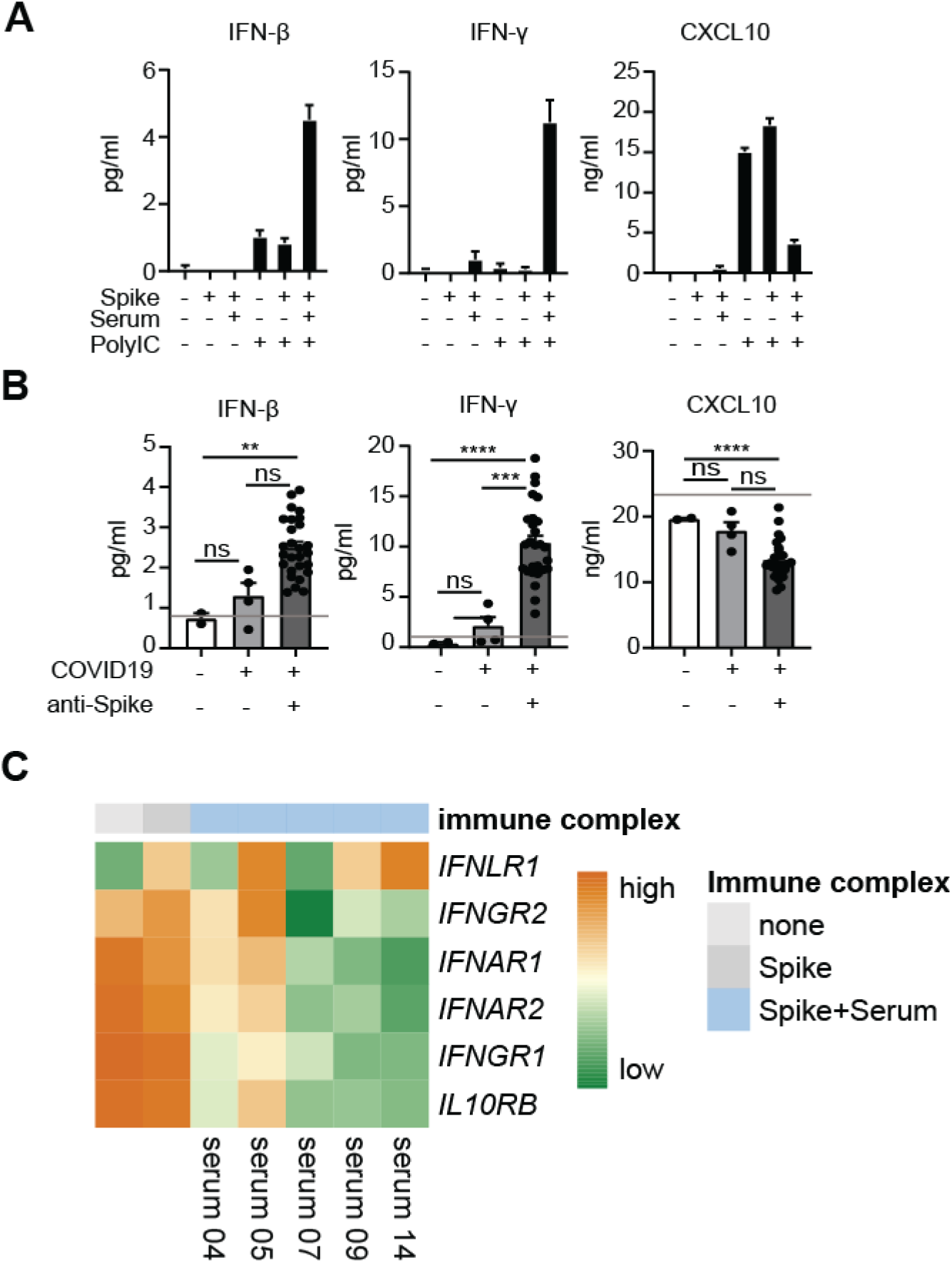
IFN responses induced by anti-Spike IgG. (A) Cytokine production by human macrophages after 6h (IFN-β and IFN-γ) or 24h (CXCL10) stimulation with (combinations of) Spike protein, COVID-19 serum, and PolyIC. Triplicate values from a representative experiment with serum (50x diluted) from five different COVID-19 patients and two different macrophage donors (mean+stdev). (B) Macrophages stimulated with Spike protein and PolyIC were co-stimulated with serum from ICU lung patients that either did not have COVID-19, had COVID-19 but were negative for anti-Spike IgG, or had COVID-19 and were positive for anti-Spike IgG. Every dot represents cytokine production after 6h (IFN-β, IFN-γ) or 24h (CXCL10) by a different serum donor (mean+SEM). Grey line indicates cytokine production induced by PolyIC-Spike. Statistics were made with Brown-Forsythe and Welch’s ANOVA test and corrected by Dunnett T3 test for multiple test correction (sample size: COVID19−: 2, COVID19+ anti-spike−: 4, COVID19+ anti-spike+: 27). Asterisks indicate significance: *\*P* < 0.05; *\*\*P* < 0.01; *\*\*\*\*P*<0.0001; ns=not significant. (C) Heatmap showing scaled log_2_ expression level of IFN receptors assessed by RNAseq upon 6h stimulation of human macrophages with PolyIC, with or without Spike protein and serum from five COVID-19 patients that tested positive for anti-Spike IgG.

**Figure S2.**
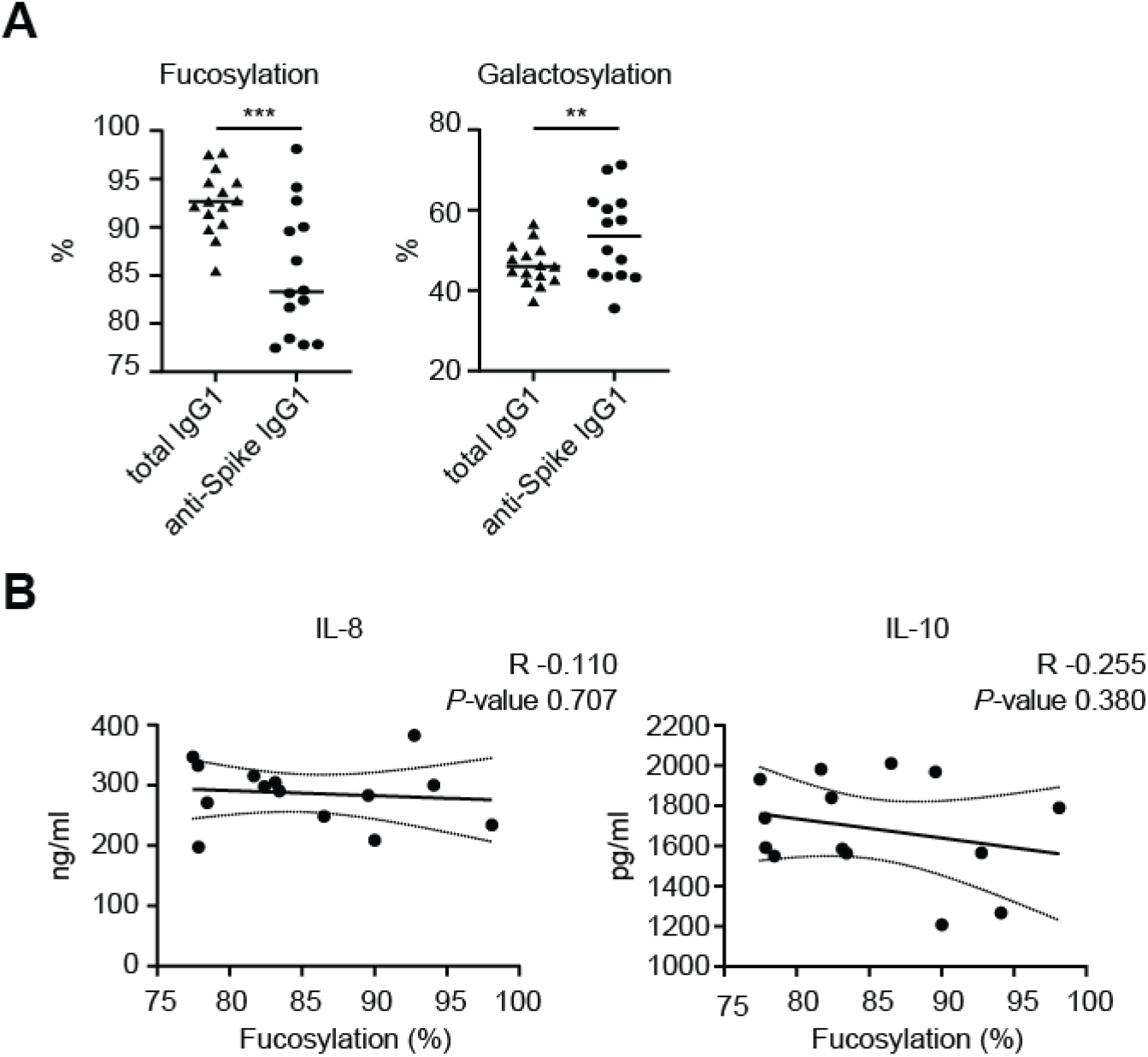
Correlation of anti-Spike IgG fucosylation and IL-8 and IL-10 production. (A) IgG1 fucosylation and galactosylation levels of total and anti-Spike specific antibodies. Statistics were calculated with a paired t test. *\*\*P* < 0.01; *\*\*\*P* <0.001. (B) Correlation graphs of fucosylation percentages of anti-Spike IgG1 from COVID-19 serum against cytokine production of macrophages after stimulation as in Fig. 3A. The Pearson correlation coefficient (R) and *P* value are illustrated on each graph with 95% confidence bands of the best-fit line.

**Figure S3.**
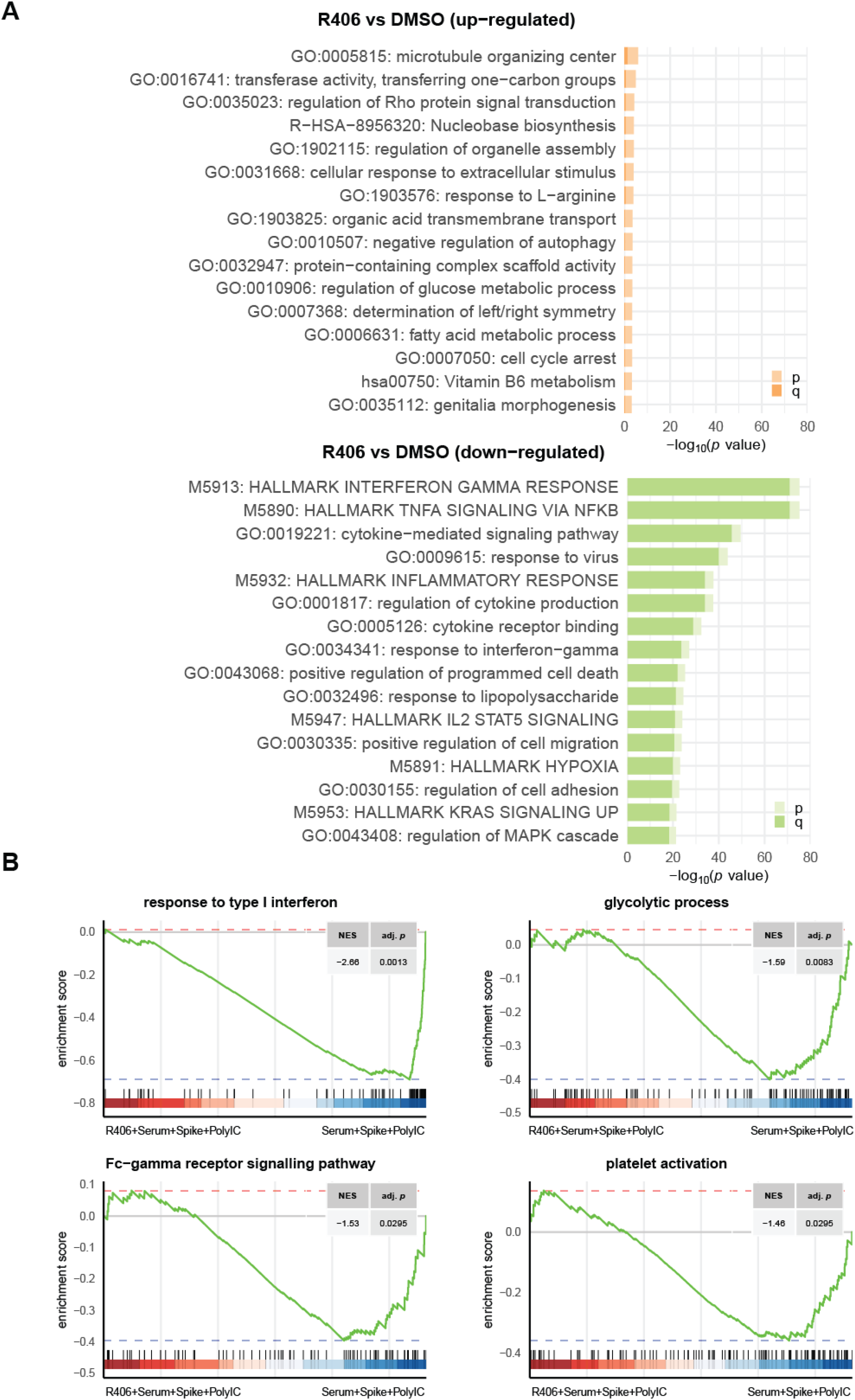
Gene regulation by R406. (A) Differentially regulated genes were defined by a false discovery rate (FDR) <0.05 and an absolute log_2_ fold-change higher than 1. Pathway enrichment analyses were performed using the Metascape on 2020-06-26. (B) Gene set enrichment analysis (GSEA) of curated gene sets suppressed by R406: Fc-gamma receptor signaling pathway (GO:0038094), glycolytic process (GO:0006096), platelet activation (GO:0030168), response to type I interferon (GO:0034340). NES stands for normalized enrichment score and adj. P represents the BH-adjusted P value.

**Table S1.** COVID-19 patient characteristics. Ventilation indicates that these patients received mechanical ventilation. The numbers in the columns of “Hospitalization” and “Sampling” indicate the days after onset of symptoms, which for some patients were unknown. IgG1 glycosylation percentages indicate the glycosylation of anti-Spike IgG1 of these patients. Donors 18 and 30 were intensive care lung disease patients that did not have COVID-19 (not included in this table). CCD: Chronic cardiac disease, including congenital heart disease (not hypertension). CPD: Chronic pulmonary disease (not asthma). CKD: Chronic kidney disease. Liver dis.: Moderate or severe liver disease. CHD: Chronic hematologic disease.

| Patient | Gender | Age | Co-morbidities | Obese | Ventilation | Outcome | Hospitalization | Sampling | IgG1 glycosylation |  |  |  |
| --- | --- | --- | --- | --- | --- | --- | --- | --- | --- | --- | --- | --- |
|  |  |  |  |  |  |  |  |  | Fucosylation % | Galactosylation % | Sialylation % | Bisection % |
| 1 | Male | 63 | Diabetes | No | Yes | recovered | 10 | 25 | 83.2 | 43.3 | 7.6 | 2.0 |
| 2 | Male | 59 | Diabetes | Yes | Yes | recovered | 7 | 21 | 81.7 | 57.6 | 9.5 | 3.3 |
| 4 | Female | 51 | None | Yes | Yes | recovered | 6 | 24 | 98.1 | 43.5 | 8.4 | 3.1 |
| 5 | Male | 70 | CHD | No | Yes | deceased | onset unknown | onset unknown | 94.1 | 35.6 | 5.8 | 4.4 |
| 6 | Male | 66 | CCD | Yes | Yes | recovered | 7 | 23 | 77.8 | 44.2 | 6.9 | 2.5 |
| 7 | Male | 61 | None | No | No | recovered | 6 | 24 | 77.5 | 50.1 | 7.8 | 8.0 |
| 8 | Male | 61 | Hypertension | No | No | recovered | 8 | 20 | 89.6 | 43.8 | 8.1 | 3.6 |
| 9 | Female | 55 | None | Yes | No | recovered | 7 | 15 | 78.5 | 60.3 | 9.2 | 2.8 |
| 10 | Female | 52 | Hypertension | Yes | Yes | recovered | 2 | 8 |  |  |  |  |
| 11 | Female | 46 | None | No | Yes | deceased | 7 | 13 | 83.4 | 71.3 | 11.6 | 3.7 |
| 13 | Male | 63 | None | No | Yes | deceased | 31 | 50 | 86.5 | 47.7 | 9.3 | 2.8 |
| 14 | Female | 50 | Hypertension | Yes | Yes | recovered | 14 | 26 | 77.8 | 56.9 | 9.1 | 3.8 |
| 15 | Male | 67 | Autoimmune dis. | No | Yes | recovered | onset unknown | onset unknown | 90.0 | 62.0 | 13.6 | 6.8 |
| 16 | Female | 71 | None | No | Yes | deceased | 5 | 19 | 82.4 | 61.7 | 10.8 | 4.4 |
| 17 | Male | 65 | CCD; diabetes; hypertension | No | Yes | recovered | 10 | 19 | 92.8 | 70.1 | 12.7 | 3.8 |
| 19 | Male | 54 | CCD | N/A | Yes | recovered | onset unknown | onset unknown |  |  |  |  |
| 21 | Male | 59 | CCD; hypertension | No | Yes | recovered | 14 | 39 |  |  |  |  |
| 22 | Male | 74 | CPD; hypertension | No | Yes | deceased | 10 | 40 |  |  |  |  |
| 23 | Male | 61 | Asthma; autoimmune dis.; diabetes | Yes | Yes | recovered | 17 | 46 |  |  |  |  |
| 24 | Male | 60 | CPD; diabetes | Yes | Yes | deceased | 4 | 26 |  |  |  |  |
| 25 | Male | 72 | Diabetes; hypertension; rheumatic dis. | No | Yes | recovered | 7 | 31 |  |  |  |  |
| 26 | Female | 79 | Autoimmun.; CCD; CKD; CPD; diabetes; hypertension | No | No | recovered | onset unknown | onset unknown |  |  |  |  |
| 27 | Female | 64 | None | Yes | No | recovered | 3 | 7 |  |  |  |  |
| 28 | Male | 61 | Hypertension; liver dis. | Yes | Yes | recovered | 7 | 12 |  |  |  |  |
| 29 | Male | 56 | Liver dis. | No | No | deceased | 9 | 14 |  |  |  |  |
| 31 | Male | 61 | CCD; diabetes; hypertension | Yes | Yes | recovered | 7 | 12 |  |  |  |  |
| 32 | Male | 66 | None | No | Yes | deceased | 7 | 20 |  |  |  |  |
| 33 | Female | 62 | CPD | No | Yes | recovered | 7 | 10 |  |  |  |  |
| 34 | Male | 59 | Diabetes | No | Yes | recovered | 4 | 5 |  |  |  |  |
| 35 | Male | 78 | CCD; CKD | No | Yes | recovered | 4 | 15 |  |  |  |  |
| 36 | Male | 46 | Autoimmun dis.; liver dis. | No | Yes | recovered | 8 | 15 |  |  |  |  |

**Table S2.** Glycosylation of COVA1-18 IgGs. Levels of fucose, galactose, sialic acid, and bisection of COVA1-18 glyco-variants was measured as described previously(*26*).

|  | **Fucosylation %** | **Galactosylation %** | **Sialylation %** | **Bisection %** |
| --- | --- | --- | --- | --- |
| COVA1-18 | 97.8 | 19.6 | 1.1 | 2.4 |
| COVA1-18 low fucose high galactose | 9.1 | 77.6 | 5.4 | 0.2 |

## References and Notes

1. R. T. Gandhi, J. B. Lynch, C. Del Rio, Mild or Moderate Covid-19. N Engl J Med, (2020).

2. M. Z. Tay, C. M. Poh, L. Renia, P. A. MacAry, L. F. P. Ng, The trinity of COVID-19: immunity, inflammation and intervention. Nat Rev Immunol 20, 363–374 (2020).

3. M. Merad, J. C. Martin, Pathological inflammation in patients with COVID-19: a key role for monocytes and macrophages. Nat Rev Immunol 20, 355–362 (2020).

4. N. Mangalmurti, C. A. Hunter, Cytokine Storms: Understanding COVID-19. Immunity, (2020).

5. D. Blanco-Melo et al., Imbalanced Host Response to SARS-CoV-2 Drives Development of COVID-19. Cell 181, 1036–1045 e1039 (2020).

6. E. J. Giamarellos-Bourboulis et al., Complex Immune Dysregulation in COVID-19 Patients with Severe Respiratory Failure. Cell Host Microbe 27, 992–1000 e1003 (2020).

7. T. Herold et al., Elevated levels of IL-6 and CRP predict the need for mechanical ventilation in COVID-19. J Allergy Clin Immunol, (2020).

8. J. Hadjadj et al., Impaired type I interferon activity and exacerbated inflammatory responses in severe Covid-19 patients. medRxiv, (2020).

9. L. Liu et al., Anti-spike IgG causes severe acute lung injury by skewing macrophage responses during acute SARS-CoV infection. JCI Insight 4, (2019).

10. Q. X. Long et al., Antibody responses to SARS-CoV-2 in patients with COVID-19. Nat Med 26, 845–848 (2020).

11. J. Qu et al., Profile of IgG and IgM antibodies against severe acute respiratory syndrome coronavirus 2 (SARS-CoV-2). Clin Infect Dis, (2020).

12. K. K. To et al., Temporal profiles of viral load in posterior oropharyngeal saliva samples and serum antibody responses during infection by SARS-CoV-2: an observational cohort study. Lancet Infect Dis 20, 565–574 (2020).

13. M. Liao et al., Single-cell landscape of bronchoalveolar immune cells in patients with COVID-19. Nat Med, (2020).

14. H. J. Chen et al., Meta-Analysis of in vitro-Differentiated Macrophages Identifies Transcriptomic Signatures That Classify Disease Macrophages in vivo. Front Immunol 10, 2887 (2019).

15. W. Hoepel, K. Golebski, C. M. van Drunen, J. den Dunnen, Active control of mucosal tolerance and inflammation by human IgA and IgG antibodies. J Allergy Clin Immunol, (2020).

16. L. T. Vogelpoel, D. L. Baeten, E. C. de Jong, J. den Dunnen, Control of cytokine production by human fc gamma receptors: implications for pathogen defense and autoimmunity. Front Immunol 6, 79 (2015).

17. C. Huang et al., Clinical features of patients infected with 2019 novel coronavirus in Wuhan, China. Lancet 395, 497–506 (2020).

18. L. Li, Q. Huang, D. C. Wang, D. H. Ingbar, X. Wang, Acute lung injury in patients with COVID-19 infection. Clin Transl Med 10, 20–27 (2020).

19. R. C. Becker, COVID-19 update: Covid-19-associated coagulopathy. J Thromb Thrombolysis, (2020).

20. L. Botros et al., Bosutinib prevents vascular leakage by reducing focal adhesion turnover and reinforcing junctional integrity. J Cell Sci 133, (2020).

21. X. D. Manz et al., In Vitro Microfluidic Disease Model to Study Whole Blood-Endothelial Interactions and Blood Clot Dynamics in Real-Time. J Vis Exp, (2020).

22. P. J. M. Brouwer et al., Potent neutralizing antibodies from COVID-19 patients define multiple targets of vulnerability. Science, (2020).

23. H. Ma et al., COVID-19 diagnosis and study of serum SARS-CoV-2 specific IgA, IgM and IgG by chemiluminescence immunoanalysis. medRxiv, (2020).

24. G. Vidarsson, G. Dekkers, T. Rispens, IgG subclasses and allotypes: from structure to effector functions. Front Immunol 5, 520 (2014).

25. M. F. Jennewein, G. Alter, The Immunoregulatory Roles of Antibody Glycosylation. Trends Immunol 38, 358–372 (2017).

26. M. D. Larsen et al., Afucosylated immunoglobulin G responses are a hallmark of enveloped virus infections and show an exacerbated phenotype in COVID-19. bioRxiv, (2020).

27. S. Chakraborty et al., Symptomatic SARS-CoV-2 infections display specific IgG Fc structures. medRxiv, (2020).

28. W. Hoepel et al., FcgammaR-TLR Cross-Talk Enhances TNF Production by Human Monocyte-Derived DCs via IRF5-Dependent Gene Transcription and Glycolytic Reprogramming. Front Immunol 10, 739 (2019).

29. L. T. Vogelpoel et al., Fc gamma receptor-TLR cross-talk elicits pro-inflammatory cytokine production by human M2 macrophages. Nat Commun 5, 5444 (2014).

30. N. T. Connell, N. Berliner, Fostamatinib for the treatment of chronic immune thrombocytopenia. Blood 133, 2027–2030 (2019).

31. M. Guilliams, P. Bruhns, Y. Saeys, H. Hammad, B. N. Lambrecht, The function of Fcgamma receptors in dendritic cells and macrophages. Nat Rev Immunol 14, 94–108 (2014).

32. D. M. Del Valle et al., An inflammatory cytokine signature helps predict COVID-19 severity and death. medRxiv, (2020).

33. M. E. Ackerman et al., Natural variation in Fc glycosylation of HIV-specific antibodies impacts antiviral activity. J Clin Invest 123, 2183–2192 (2013).

34. T. T. Wang et al., IgG antibodies to dengue enhanced for FcgammaRIIIA binding determine disease severity. Science 355, 395–398 (2017).

35. S. B. Halstead, Dengue Antibody-Dependent Enhancement: Knowns and Unknowns. Microbiol Spectr 2, (2014).

36. T. Zohar, G. Alter, Dissecting antibody-mediated protection against SARS-CoV-2. Nat Rev Immunol, (2020).

37. K. Golebski et al., FcgammaRIII stimulation breaks the tolerance of human nasal epithelial cells to bacteria through cross-talk with TLR4. Mucosal Immunol 12, 425–433 (2019).

38. R. L. Chua et al., COVID-19 severity correlates with airway epithelium-immune cell interactions identified by single-cell analysis. Nat Biotechnol, (2020).

39. M. Alimova et al., A High Content Screen for Mucin-1-Reducing Compounds Identifies Fostamatinib as a Candidate for Rapid Repurposing for Acute Lung Injury during the COVID-19 pandemic. bioRxiv, (2020).

40. J. Banga et al., Inhibition of IRF5 cellular activity with cell-penetrating peptides that target homodimerization. Sci Adv 6, eaay1057 (2020).

41. B. Shen et al., Proteomic and Metabolomic Characterization of COVID-19 Patient Sera. Cell, (2020).

42. G. Dekkers et al., Multi-level glyco-engineering techniques to generate IgG with defined Fc-glycans. Sci Rep 6, 36964 (2016).

43. M. W. J. Smeets, M. J. Mourik, H. W. M. Niessen, P. L. Hordijk, Stasis Promotes Erythrocyte Adhesion to von Willebrand Factor. Arterioscler Thromb Vasc Biol 37, 1618–1627 (2017).

44. A. Dobin et al., STAR: ultrafast universal RNA-seq aligner. Bioinformatics 29, 15–21 (2013).

45. H. Li et al., The Sequence Alignment/Map format and SAMtools. Bioinformatics 25, 2078–2079 (2009).

46. M. I. Love, W. Huber, S. Anders, Moderated estimation of fold change and dispersion for RNA-seq data with DESeq2. Genome Biol 15, 550 (2014).

47. D. J. McCarthy, Y. Chen, G. K. Smyth, Differential expression analysis of multifactor RNA-Seq experiments with respect to biological variation. Nucleic Acids Res 40, 4288–4297 (2012).

48. Y. Zhou et al., Metascape provides a biologist-oriented resource for the analysis of systems-level datasets. Nat Commun 10, 1523 (2019).

49. G. Korotkevich, V. Sukhov, A. Sergushichev, Fast gene set enrichment analysis. bioRxiv, (2019).

50. S. Durinck, P. T. Spellman, E. Birney, W. Huber, Mapping identifiers for the integration of genomic datasets with the R/Bioconductor package biomaRt. Nat Protoc 4, 1184–1191 (2009).

